# A novel plasmid-based experimental system in *Saccharomyces cerevisiae* that enables the introduction of 10 different plasmids into cells

**DOI:** 10.1101/2023.05.01.538886

**Authors:** Geyao Dong, Tsuyoshi Nakai, Tetsuo Matsuzaki

## Abstract

The budding yeast *Saccharomyces cerevisiae* is commonly used as an expression platform to produce valuable compounds. Yeast-based genetics research can uniquely utilize use auxotrophy in transformant selection: auxotrophic complementation by an auxotrophic marker gene on exogenous DNA (such as plasmids). However, the number of auxotrophic nutrients required restricts the number of plasmids maintained by the cells. We therefore developed novel Δ10 strains that are auxotrophic for 10 different nutrients, and new plasmids with two multicloning sites and auxotrophic markers for use in Δ10 strains. We confirmed that Δ10 strains could maintain 10 types of these plasmids. Because each plasmid can express two different genes, this Δ10 strain-based expression system has the potential to co-express a maximum of 20 different proteins.

## Introduction

The budding yeast *Saccharomyces cerevisiae* is one of the most widely used eukaryotic model organisms in laboratory experiments, due to the ease and precision with which its genome can be manipulated and the high degree of conservation of cellular processes and molecular pathways with higher eukaryotes, including humans [1–4]. Besides being an important model organism for basic studies, *S. cerevisiae* serves as an invaluable tool in the field of biotechnology [5,6]. Recent developments in yeast synthetic biology have facilitated the engineering of yeast to produce fuels, food ingredients, and biopharmaceuticals [7–15].

An important aspect of yeast genetics is the use of multiply auxotrophic yeast strains: strains that carry mutations in genes involved in biosynthetic pathways, typically genes essential for amino acid or nucleotide biosynthesis [16,17]. An example is the *URA3* gene, which encodes orotidine-5’-phosphate decarboxylase, an essential enzyme in the de novo biosynthesis of pyrimidine [18]. *URA3*null mutants are defective in uracil biosynthesis and therefore cannot grow on media lacking uracil. With such strains, transformants containing exogenous DNA (such as plasmids) encoding wild-type *URA3* can be selected as viable clones on media lacking uracil. Thus, *URA*3 is used as an auxotrophic marker for *URA3* null mutants. Similarly, *HIS3*, *LEU2*, *TRP1*, *LYS2*, *MET15*, and *ADE2* encode essential enzymes for L-histidine, L-leucine, L-tryptophan, L-lysine, L-methionine, and adenine biosynthesis, respectively, and these genes are often used as auxotrophic markers [16,17].

W303 and BY4741/BY4742 are widely used laboratory strains [16,17]. W303 carries mutations in*ADE2*, *TRP1*, *LEU2*, and *HIS3*, and these genes are also frequently employed auxotrophic markers. Meanwhile, BY strains are wild-type for *TRP1* and *ADE2*, but BY4741 and BY4742 are deficient in *MET15* and *LYS2*, respectively. The limited availability of auxotrophic markers restricts gene manipulation by auxotrophy. This complicates efforts to engineer yeast to produce chemicals, because a large number of heterologous genes must be introduced to reconstitute the biosynthesis pathways of the target products. For example, the total synthesis of opioids in yeast has been achieved by co-expressing more than 20 heterologous genes [19].

In this study, we created novel Δ10 strains (YMT183‒185) that carry null mutations in 10 different auxotrophic markers, and a series of plasmids that encode a wild-type allele of an auxotrophic markerdeleted in Δ10 strains and also contain two multi-cloning sites. We also confirmed that 10 plasmids could be introduced into Δ10. Because two different genes can be co-expressed from a single plasmid, this study may make it feasible to co-express a maximum of 20 different genes.

## Materials and methods

### 2.1. Strains and plasmids

S288C, BY4741, BY4742, and BY4743 were purchased from Open Biosystems. The *Δtrp1*, *Δade2*,*Δthr1*, *Δarg1*, and *Δtyr1* strains, which were used for constructing Δ10 strains YMT53‒YMT55 (Fig. 1), were from a collection of nonessential gene deletion strains purchased from Open Biosystems. pESC-URA, pESC-LEU, pESC-HIS, and pESC-TRP were purchased from Agilent Technologies.

### 2.2. Construction of Δ10 strains

A schematic diagram for the construction of Δ10 strains (YMT53–YMT55) is shown in Fig. 1A. First, BY4743 was sporulated and the resulting tetrads were dissected. Colonies that did not grow on media lacking either lysine or methionine (i.e., these colonies were auxotrophic for lysine and methionine, respectively) were picked and crossed to *Δtrp1* or *Δade2* cells. Tetrads resulting from a cross to *Δtrp1* and *Δade2* cells were dissected, yielding colonies auxotrophic for tryptophan and adenine, respectively. The obtained strains were crossed with each other and the resulting diploids were sporulated. After tetrad dissection, we obtained YMT43 and YMT44 colonies auxotrophic for both tryptophan and adenine (YMT43: *MAT*a *ura3Δ0 leu2Δ0 his3Δ1 met15Δ0 lys2Δ0 trp1Δ::KanMX ade2Δ::KanMX*; YMT44: *MAT*α *ura3Δ0 leu2Δ0 his3Δ1 met15Δ0 lys2Δ0 trp1Δ::KanMX ade2Δ::KanMX*).

**Fig. 1.**
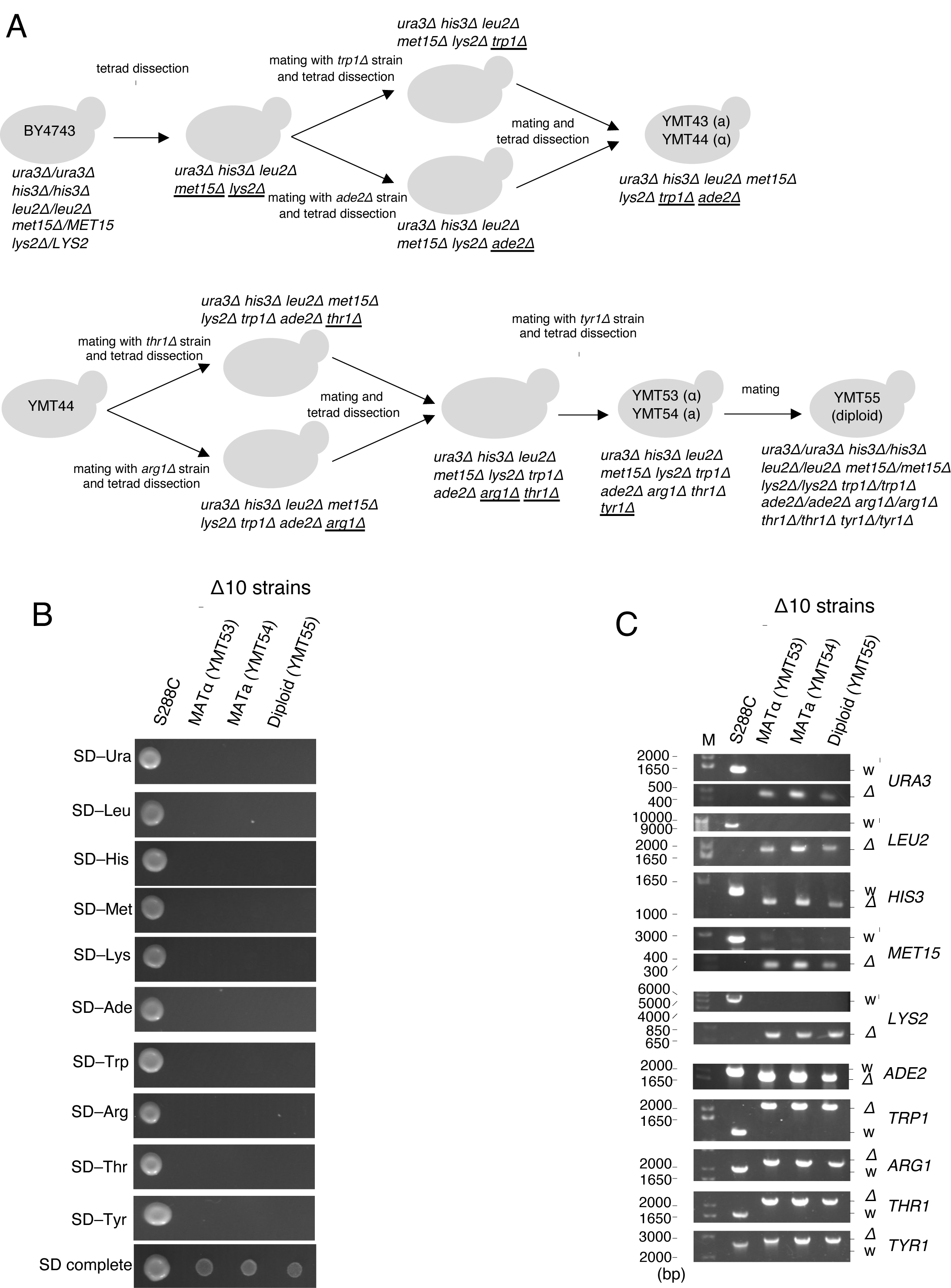
Construction of Δ10 strains (A) Schematic procedure for construction of the Δ10 strains YMT53–YMT55. Underlined letters indicate the mutation introduced at each step. (B) Deletion of each auxotrophic marker in Δ10 strains was confirmed by PCR. M: 10-kbp DNA marker. (C) Δ10 strains were auxotrophic for 10 different nutrients: uracil, leucine, histidine, methionine, lysine, adenine, tryptophan, arginine, threonine, and tyrosine. Cells were spotted and cultured on indicated plates for 2 days at 27°C. The S288C strain, which does not exhibit any auxotrophy and is the ancestor of BY4741/BY4742, was used as a control.

YMT44 was crossed to *Δthr1* or *Δarg1* cells and the resulting diploids were sporulated. Spores with the following genotypes were selected: *MAT*a *ura3Δ0 leu2Δ0 his3Δ1 met15Δ0 lys2Δ0 trp1Δ::KanMX ade2Δ::KanMX thr1Δ::KanMX* (from a cross between YMT44 and *Δthr1*) and *MAT*α *ura3Δ0 leu2Δ0 his3Δ1 met15Δ0 lys2Δ0 trp1Δ::KanMX ade2Δ::KanMX arg1Δ::KanMX* (from a cross between YMT44 and *Δarg1*). These cells were crossed with each other and the resulting diploids were sporulated. After tetrad dissection, we obtained a strain auxotrophic for both threonine and arginine (*MAT*α *ura3Δ0 leu2Δ0 his3Δ1 met15Δ0 lys2Δ0 trp1Δ::KanMX ade2Δ::KanMX thr1Δ::KanMX arg1Δ::KanMX*). This strain was crossed to *tyr1Δ* cells and the resulting diploids were sporulated. Spores auxotrophic for lysine, tryptophan, adenine, threonine, arginine, and tyrosine were selected, obtaining YMT53 (*MAT*α) and YMT54 (*MAT*a). Genotypes of YMT53 and YMT54 are as follows: YMT53: *MAT*α *ura3Δ0 leu2Δ0 his3Δ1 met15Δ0 lys2Δ0 trp1Δ::KanMX ade2Δ::KanMX thr1Δ::KanMX arg1Δ::KanMX tyr1Δ::KanMX*; YMT54: *MAT*a *ura3Δ0 leu2Δ0 his3Δ1 met15Δ0 lys2Δ0 trp1Δ::KanMX ade2Δ::KanMX thr1Δ::KanMX arg1Δ::KanMX tyr1Δ::KanMX*. YMT55 was constructed by a cross between YMT53 and YMT54.

A schematic diagram of *KanMX* removal from auxotrophic marker gene loci in YMT53 and YMY54 is shown in Fig. 2A (see also Supplemental Figure 1).

**Fig. 2.**
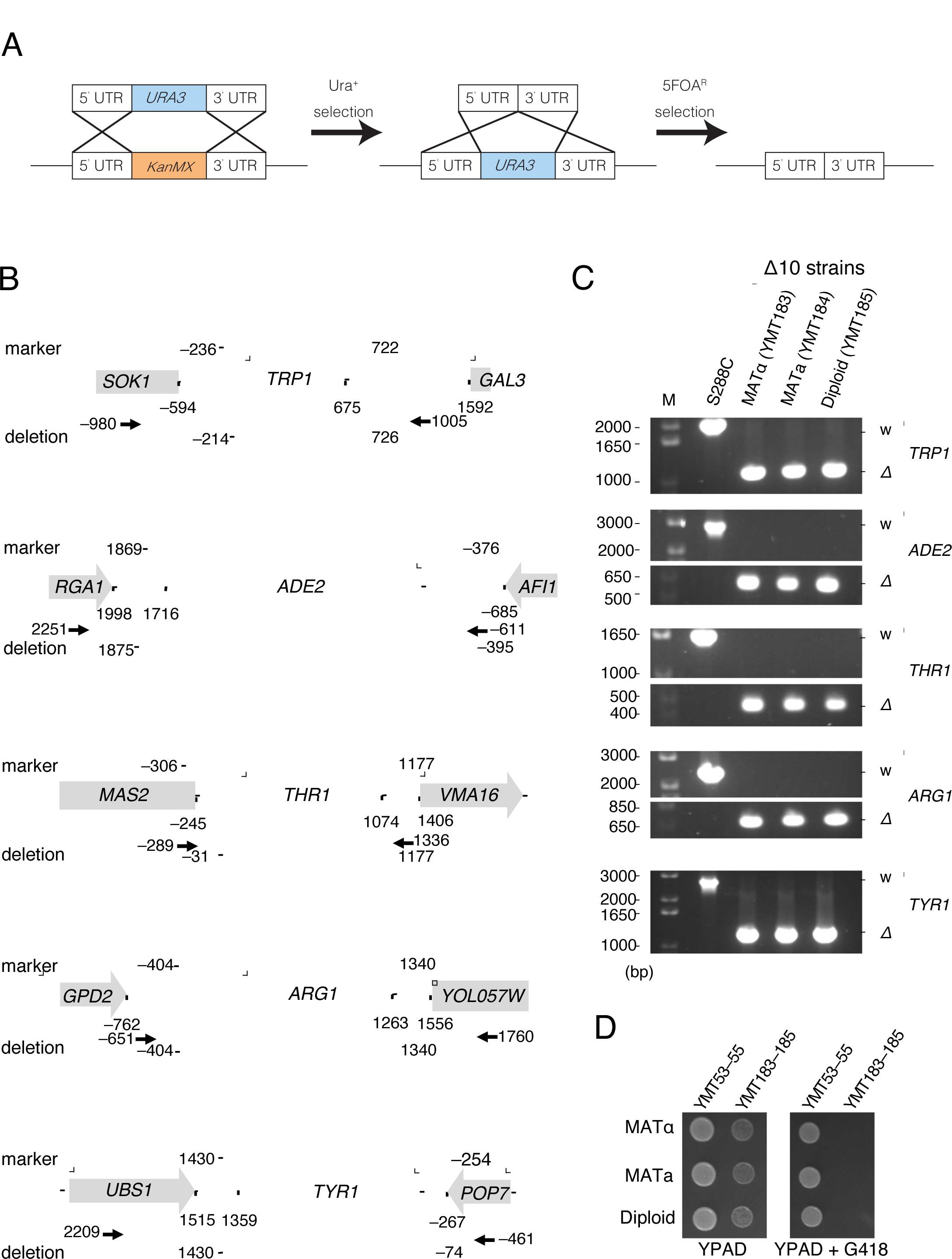
Removal of antibiotic markers (A) Method of removing the antibiotic marker from a target locus. A DNA fragment of the *URA3* ORF flanked by upstream and downstream regions of the target auxotrophic marker gene was transformed into 1¢10 cells, and cells were grown on SD−Ura plates. The resulting colonies on SD−Ura (Ura+ cells) were selected. Ura+ cells were further transformed by a DNA fragment that contained upstream and downstream regions of the target auxotrophic marker gene, then grown on SD+5-FOA plates. The resulting colonies on SD+5-FOA (FOA^R^ cells) were selected; the cells were expected to have lost the antibiotic marker from the target locus. (B) Deleted regions in each auxotrophic marker and overlap with vector segments. The genomic region encompassing each auxotrophic marker gene is presented. The definitions of nucleotide numbers were based on assigning +1 to the A of the first ATG in the auxotrophic marker ORF. Lines above each marker locus indicate deleted regions. Lines under each marker locus indicate genomic regions that are cloned into the corresponding pESC vector (Fig. 3B). Arrows indicate primer sets used in PCR to confirm deletions in each locus (Fig. 2C), and the nucleotide numbers corresponding to the 5’-nucleotide of each primer are shown. (C) Removal of the antibiotic marker was confirmed by PCR. M: 10-kbp DNA marker. (D) The new 1¢10 strains, YMT183–YMT185, were sensitive to G418. Cells were spotted and cultured on YPAD or YPAD+G418 plates for 1 day at 27°C.

For removal of *KanMX* from the *ADE2* locus, regions upstream and downstream of the *ADE2* open reading frame (ORF) were PCR-amplified using the M29/M413 and M30/M414 primer sets, respectively. Then resultant PCR DNA fragments and the primer set M29/M30 were used for second PCR, resulting in DNA fragments containing *URA3* flanked by regions upstream and downstream of the *ADE2* ORF. These DNA fragments were transformed into yeast and cells were grown on SD−Ura plates, resulting in cells that carried the *ade2Δ::URA3* allele. To remove integrated *URA3* from the *ADE2* locus, regions upstream and downstream of the *ADE2* ORF were PCR-amplified using the M29/M441 and M30/M440 primer sets, respectively. A second PCR was performed using the obtained upstream and downstream *ADE2* regions as templates and the primer set M29/M30, and this resulted in DNA fragments containing upstream and downstream *ADE2* flanking regions. The obtained DNA fragments were transformed into yeast and cells were grown on SD media containing 5-fluoro-orotic acid (5-FOA), resulting in cells that carried the *ade2Δ4* allele.

Removal of *KanMX* from *TYR1*, *TRP1*, *ARG1*, and *THR1* loci was performed in similar ways. For the construction of DNA fragments containing the *URA3* ORF, first PCRs were performed using the following primer sets: *TYR1*, M36/M437 and M402/M438; *TRP1*, M333/M422 and M334/M423; *ARG1*, M40/M395 and M396/M411; and *THR1*, M38/M393 and M39/M390. Then, second PCRs were performed using the respective PCR-amplified DNA fragments and the following primer sets: *TYR1*, M36/M402; *TRP1*, M333/M334; *ARG1*, M40/M411; and *THR1*, M38/M39. For construction of DNA fragments used to remove *URA3* from the target locus, first PCRs were performed using the following primer sets: *TYR1*, M36/M470 and M37/M469; *TRP1*, M433/M435 and M334/M434; *ARG1*, M40/M468 and M401/M467; and *THR1*, M38/M487 and M39/M486. Then, second PCRs were performed using the respective PCR amplified DNA fragments and following primer sets: *TYR1*, M36/M37; *TRP1*, M334 /M433; *ARG1*, M40/M401; and *THR1*, M38/M39.

The resulting YMT183 and YMT184 strains (from YMT53 and YMT54, respectively) were crossed with each other to produce a diploid strain, YMT185.

### 2.3. Construction of pESC plasmids with new markers

For the construction of pESC-MET^Pro^, pESC-ADE^Pro^, and pESC-THR^Pro^, PCRs were performed using the yeast genome as a template DNA and the M34/M35, M29/M33, and M38/M39 primer sets, respectively. The amplified fragments were cloned into the PstI and StuI sites of pESC-URA. For construction of pESC-LYS^Pro^, pESC-TYR^Pro^, and pESC-ARG^Pro^, PCRs were performed using the yeast genome as a template DNA and the M31/M32, M36/M37, and M40/M41 primer sets, respectively. The amplified fragments were cloned into the NdeI and StuI sites of pESC-URA.

For the construction of pESC-THR, the NheI site present in the *THR1* marker on pESC-THR^Pro^ was removed by site-directed mutagenesis using the M425/M426 primer sets. The remaining *URA3* ORF and promoter sequences were removed by inverse PCR using the M446/M448 and M38/442 primer sets, respectively.

For the construction of pESC-ADE, the BglII site present in the *ADE2* marker on pESC-ADE^Pro^ was removed by site-directed mutagenesis using the M405/M406 primer sets. The remaining *URA3* ORF and promoter sequences were removed by inverse PCR using the M448/M460 and M442/455 primer sets, respectively.

For the construction of pESC-ARG, the KpnI and ClaI sites present in the *ARG1* marker on pESC-ARG^Pro^ were removed by site-directed mutagenesis using the M451/M452 and M456/M457 primer sets, respectively. The remaining *URA3* ORF and promoter sequences were removed by inverse PCR using the M41/M448 and M442/461 primer sets, respectively.

For the construction of pESC-MET, the EcoRI site present in the *MET15* marker on pESC-MET^Pro^ was removed by site-directed mutagenesis using the M418/M419 primer sets. The remaining *URA3* ORF and promoter sequences were removed by inverse PCR using the M448/M464 and M442/462 primer sets, respectively.

For the construction of pESC-LYS, the BamHI, BglII, and XhoI sites present in the *LYS2* marker on pESC-LYS^Pro^ were removed by site-directed mutagenesis using the of M407/M408, M416/M417, and M429/M430 primer sets, respectively. The remaining *URA3* ORF and promoter sequences were removed by inverse PCR using the M32/M448 and M442/M443 primer sets, respectively.

For the construction of pESC-TYR, the EcoRI, HindIII, SacI, and SpeI sites present in the *TYR1* marker on pESC-TYR^Pro^ were removed by site-directed mutagenesis using the M431/M432, M444/M445, M453/M454, and M458/M459 primer sets, respectively. The remaining *URA3* ORF and promoter sequences were removed by inverse PCR using the M448/M473 and M442/M474 primer sets, respectively.

### 2.4. PCR analysis

To confirm the deletion alleles of auxotrophic marker genes in Δ10 strains (YMT53–YMT55), PCRs were performed using the genome of each strain as a template, along with primer sets flanking the target gene. The primer sets were as follows: *URA3*, M335/M336; *LEU2*, M330/M331; *HIS3*, M328/M329; *MET15*, M34/M332; *TRP1*, M333/M334; *ADE2*, M327/M356; *LYS2*, M31/M32; *ARG1*, M40/M41; *THR1*, M38/M39; and *TYR1*, M36/M37.

To confirm the *trp1Δ4*, *ade2Δ4*, *thr1Δ2*, *arg1Δ0*, and *tyr1Δ0* alleles in Δ10 strains (YMT183– YMT185), PCRs were performed in similar ways. The primer sets were as follows: *TRP1*, M334/M433; *ADE2*, M29/M33; *THR1*, M409/M410; *ARG1*, M40/M401; and *TYR1*, M42/M402.

To confirm that Δ10 cells could maintain 10 types of pESC plasmids, PCRs were performed using DNA extracted from Δ10 cells transformed by these plasmids as a template, along with primer sets specific for marker segments on the plasmids. The primer sets were as follows: *URA3*, M354/M490; *LEU2*, M490/M525; *HIS3*, M350/M490; *TRP1*, M352/M490; *ADE2*, M460/M490; *MET15*, M419/M490; *LYS2*, M32/M490; *ARG1*, M452/M490; *THR1*, M426/M490; and *TYR1*, M431/M490.

## Results

### 3.1. Construction of Δ10 strains

Using the Kyoto Encyclopedia of Genes and Genomes (KEGG) database and the Saccharomyces Genome Database (SGD), we searched for auxotrophic markers that are not used in the W303 and BY4741/BY4742 strains [20,21]. We identified the candidate genes *THR1*, *ARG1*, and *TYR1*, which encode enzymes essential for the biosynthesis of L-threonine, L-arginine, and L-tyrosine, respectively[22]. Starting with BY4743 [17], we aimed to create strains that carried mutations in a total of 10auxotrophic marker genes: *URA3*, *LEU2*, *HIS3*, *MET15*, *TRP1*, *ADE2*, *LYS2*, *THR1*, *ARG1*, and *TYR1*. Fig. 1A shows a schematic of the strategy. First, BY4743 cells were sporulated under starvation conditions, and the resulting tetrads were segregated. Because BY4743 carries heterozygous deletions in *MET15* and *LYS2*, a strain that carries homozygous deletions in these two genes was obtained by selecting segregated cells that did not grow on either lysine- or methionine-deficient media. This strain was further crossed to two strains, one with *TPR1* deletion and the other with *ADE2* deletion, and the resulting diploid cells were sporulated and segregated. *URA3*, *LEU2*, *HIS3*, *MET15*, *LYS2*, and *TRP1* deletion strains and *URA3*, *LEU2*, *HIS3*, *MET15*, *LYS2*, and *ADE2* deletion strains were obtained by auxotrophic selection. By crossing these two strains, strains that carried *URA3*, *LEU2*, *HIS3*, *MET15*, *LYS2*, *TRP1*, and *ADE2* deletions (YMT43/YMT44) were obtained in a similar manner.

Next, *THR1* and *ARG1* deletion alleles were introduced into YMT44 by crossing YMT44 to *thr1Δ* cells and *arg1Δ cells*, respectively. These cells were crossed to each other, and the resulting diploids were segregated, yielding a strain that carried both *THR1* and *ARG1* deletions. This strain was further crossed to the *TYR1* deletion strain and segregated. Among the segregated cells, we obtained Δ10 strains (YMT53/YMT54) strains that exhibited auxotrophy for uracil, leucine, histidine, methionine, lysine, tryptophan, adenine, threonine, arginine, and tyrosine (Fig. 1A and 1B). The diploid Δ10 strain (YMT55) was obtained by a cross between YMT53 and YMT54. Deletions in auxotrophic markers were confirmed by PCR (Fig. 1C).

Because *TRP1*, *ADE2*, *THR1*, *ARG1*, and *TYR1* deletion alleles contained the antibiotic marker cassette *KanMX*, which confers resistance to geneticin (G418), we removed the antibiotic marker from each locus by URA3/5-FOA counterselection [23]. To exchange *KanMX* with *URA3*, Δ10 cells were transformed by a DNA fragment containing *URA3* flanked by 5’- and 3’-untranslated regions (UTRs) of the target gene. Ura+ colonies were selected on uracil-deficient media. To remove *URA3* from the target locus, Ura+ cells were further transformed by a DNA fragment containing 5’- and 3’-UTRs of the target gene. Ura-colonies were selected on media containing 5-FOA (Fig. 2A). In the latter transformation, several hundred base pairs (bp) of 5’- and 3’-UTRs were removed from the target locus so that there was no homology between the marker segment in the commonly used plasmid and the deletion in the chromosome (Fig. 2B). By repeating this, we removed the antibiotic markers from the *TRP1*, *ADE2*, *THR1*, *ARG1*, and *TYR1* loci, obtaining YMT183 and YMT184 (Fig. 2C). These modified deletion alleles of *trp1*, *ade2*, *thr1*, *arg1*, and *tyr1* are designated as *trp1Δ4*, *ade2Δ4*, *thr1Δ2*, *arg1Δ0*, and *tyr1Δ0*, respectively. The diploid strain YMT185 was obtained by a cross between YMT183 and YMT184. We confirmed that YMT183‒YMT185 no longer exhibited G418 resistance(Fig. 2D).

### 3.2 Construction of pESC plasmids with new markers

pESC plasmids are yeast expression vectors that contain two multi-cloning sites, which enables the co-expression of two genes (http://www.chem-agilent.com). Commercially available pESC-URA, pESC-LEU, pESC-HIS, and pESC-TRP plasmids contain *URA3*, *LEU2*, *HIS3*, and *TRP1* as selectable markers, respectively. We constructed new pESC plasmids for use with Δ10 strains by replacing the marker segment of pESC-URA, as follows: pESC-MET^Pro^ (*MET15* marker), pESC-ADE^Pro^ (*ADE2* marker), pESC-THR^Pro^ (*THR1* marker), pESC-LYS^Pro^ (*LYS2* marker), pESC-TYR^Pro^ (*TYR1* marker), and pESC-ARG^Pro^ (*ARG1* marker) (“Pro” denotes a prototype, because these plasmids were modified as described below). Transformation of Δ10 cells by these plasmids yielded colonies on corresponding selective media (Fig. 3A).

**Fig. 3.**
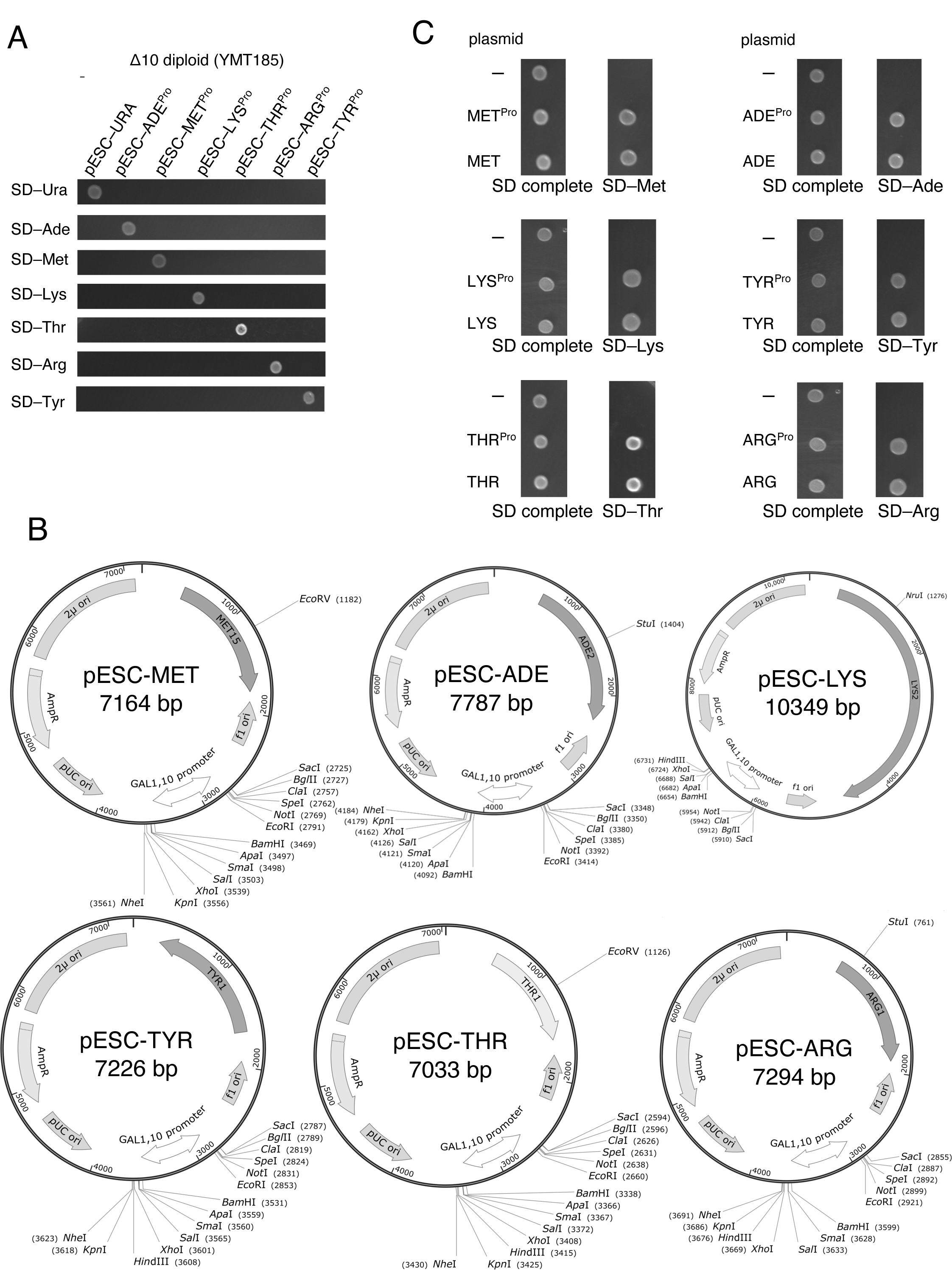
Construction of pESC plasmids with new marker genes (A) Cells carrying a pESC plasmid do not exhibit auxotrophy for the nutrient corresponding to the auxotrophic marker gene on the plasmid. Cells were spotted and cultured on the indicated plates for 2 days at 27°C. (B) Maps of new pESC plasmids. Unique restriction sites are shown. (C) Confirmation that modified pESC plasmids can be used for auxotrophic selection. Cells were spotted and cultured on the indicated plates for 2 days at 27°C.

To increase the number of unique restriction sites in the multicloning site, site-directed silent mutations were introduced into the auxotrophic markers. Specifically, the following were destroyed: the NheI site (+1054) of *THR1*; BglII site (+592) of *ADE2*; KpnI (+269) and ClaI (+806) sites of *ARG1*; EcoRI site (+415) of *MET15*; BglII (+387), XhoI (+2864), and BamHI sites (+3246) of *LYS2*; and HindIII (– 105), SpeI (+429), EcoRI (+963), and SacI sites (+1192) of *TYR1*. As a result, we obtained modified pESC plasmids: pESC-MET, pESC-ADE, pESC-LYS, pESC-TYR, pESC-THR, and pESC-ARG (Fig. 3B). Transformants with these modified pESC plasmids grow on corresponding selective media, like prototype pESC plasmids (Fig. 3C).

### 3.3. Establishment of a novel yeast protein expression system that enables simultaneous expression of a maximum of 20 different genes

We next examined whether Δ10 cells could carry all 10 plasmids. We used the plasmids to sequentially transform Δ10 cells (YMT185), and successfully obtained cells that grew on media lacking the corresponding amino acids and nucleotides (Fig. 4A, SD–ULHWAMKTRY). PCRs were performed using DNA extracted from cells that grow on SD–ULHWAMKTRY as a template, along with primer sets specific for the marker segment on each plasmid. PCR products of the expected size were amplified, demonstrating that cells grown on SD–ULHWAMKTRY carried 10 plasmids (Fig. 4B). Because each plasmid enables the co-expression of two genes, this strain has the potential to co-express a maximum of 20 different genes.

**Fig. 4.**
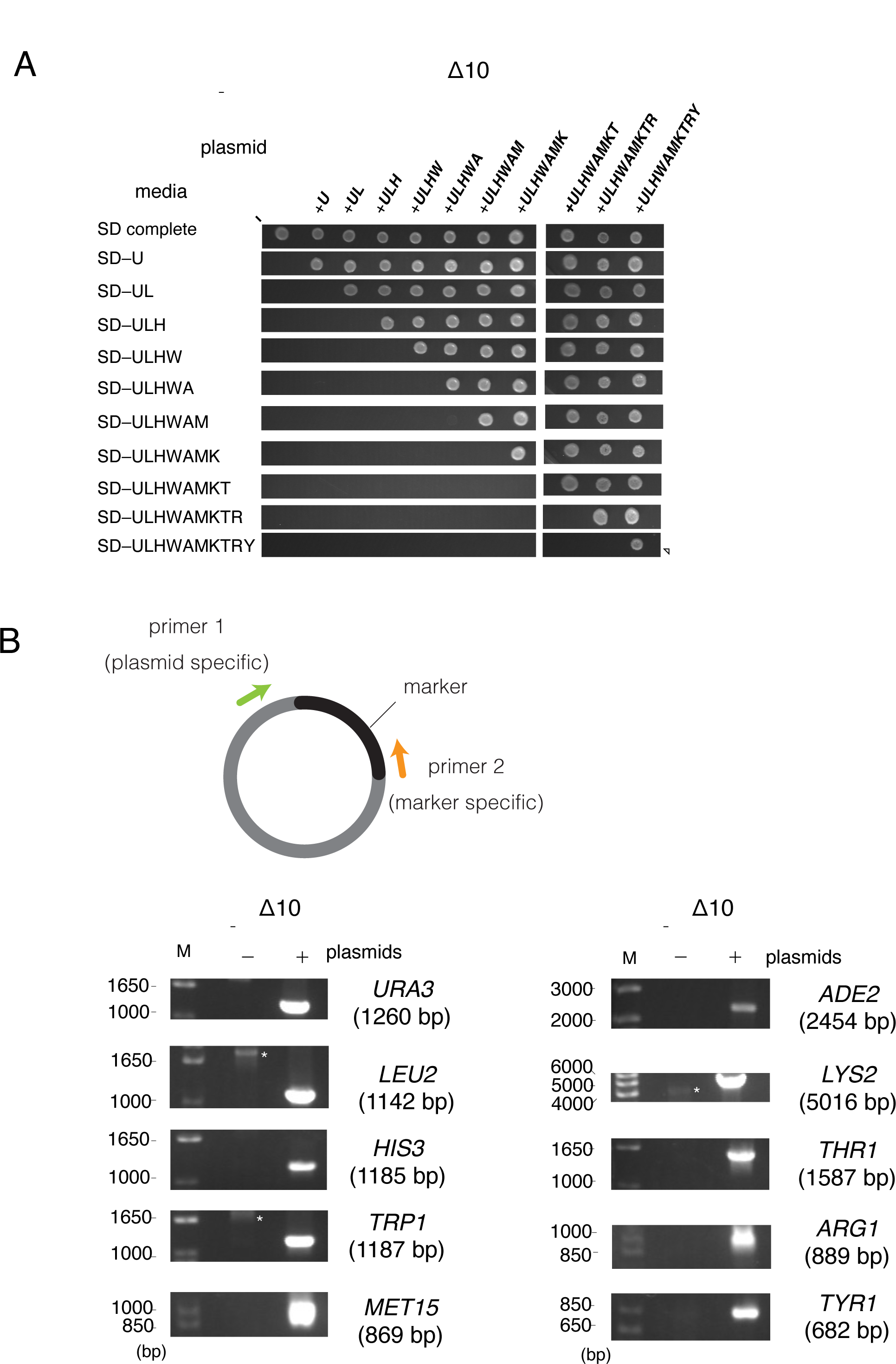
Confirmation that Δ10 cells can carry 10 pESC plasmids (A) Sequential introduction of pESC plasmids into 1¢10 cells. 1¢10 cells (YMT185) were serially transformed by pESC plasmids. Transformed cells were spotted and cultured on the indicated plates for 2 days at 27°C. pESC plasmids are indicated as follows: *U*=pESC-URA, *L*=pESC-LEU, *H*=pESC-HIS, *W*=pESC-TRP, *A*=pESC-ADE, *M*=pESC-MET, *K*=pESC-LYS, *T*=pESC-THR, *R*=pESC-ARG, *Y*=pESC-TYR. The arrowheads represent cells harboring 10 pESC plasmids. (B) Confirmation that cells grown on SD−ULHWAMKTRY harbor all 10 pESC plasmids. PCRs were performed using DNA extracted from YMT185 or cells that grow on SD–ULHWAMKTRY as a template, along with primer sets specific for the marker segment on the plasmid. Expected sizes of PCR products are indicated in each electrogram. M: 10-kbp DNA marker.

## Discussion

In this study, we established a novel protein expression system in *S. cerevisiae* that enables co-expression of a maximum of 20 genes. The commonly used yeast strains BY4741 and BY4742 carry *ura3Δ0*, *leu2Δ0*, and *his3Δ1*; in addition, the former carries *met15Δ0* while the latter carries *lys21¢0* [17]. Another widely used strain, W303, carries *leu2-3,112*, *trp1-1*, *ura3-1*, *ade2-1*, and *his3-11,15* as auxotrophic markers [16]. Due to this limited number of markers, the maximum numbers of plasmids that BY4741/BY4742 and W303 can maintain by auxotrophic selection are four and five, respectively.Thus, the Δ10-based expression system developed in this study greatly expanded the number of plasmids that can be maintained in cells by auxotrophic selection.

Besides being a suitable model organism in the field of basic biology, budding yeast is utilized as a cell factory for producing high-value compounds such as fuels, food ingredients, and biopharmaceuticals [12,13,15,24]. One of the most successful examples is human insulin, an essential biopharmaceutical in treating diabetes [25]. Glucagon, human serum albumin, and hepatitis B virus vaccine are other biopharmaceuticals produced by budding yeast (reviewed in [8,24]). Recently, the biopharmaceuticals produced by budding yeast have expanded beyond recombinant peptides and proteins. *Awan et al*. engineered budding yeast to produce and secrete benzylpenicillin, a ý-lactam antibiotic, by introducing five genes of *Penicillium chrysogenum* that are involved in benzylpenicillin biosynthesis [26]. They confirmed that spent culture media from the engineered yeast inhibited the growth of the pathogenic bacteria *Streptococcus pyogenes*, demonstrating that the yeast secreted bioactive benzylpenicillin. Galanie *et al.* established the biosynthesis of the opioid compounds thebaine and hydrocodone in budding yeast [19]. In their study, large DNA fragments encoding genes involved in opioid biosynthesis (more than 20 genes in total) were integrated into the yeast genome to reconstitute the opioid biosynthesis pathway. However, the construction of such large DNA fragments and their subsequent integration into the yeast genome are technically difficult. In contrast, the multi-expression system constructed in this study is easy to use because this system is based on the classic introduction of plasmids into yeast cells by auxotrophic complementation. Though sequential introduction of plasmids is a time-consuming process, we believe that the novel multi-expression system established here will be a useful platform for yeast cell factories.

For technical reasons, we did not investigate the copy numbers of all 10 plasmids in the Δ10 cells in this study. Two-micron plasmids, which include pESC plasmids, generally maintain 40–60 copies per cell [27]. It was recently reported that the selection marker used strongly affects the plasmid copynumber [28]. Therefore, there may be significant variation in the copy numbers of the plasmids maintained in cells. In yeast cell factories, the expression levels of heterologous genes are important determinants of the titers of the collected target compounds [24,29]. For example, in the study by*Awan et al*. mentioned above, optimizing plasmid copy numbers and the promoters of genes in the penicillin biosynthetic pathway resulted in a 30-fold increase in the yields of secreted benzylpenicillin compared to the prototype strain [26]. The current Δ10-based multi-expression system uses 2-micron plasmids in which expression of the encoded genes is under the control of the *GAL1*/*GAL10* promoters [30]. Other plasmids such as centromeric plasmids, which maintain low copy numbers per cell (one to four copies per cell) [7], and plasmids containing different promoters will be beneficial to regulate the expression levels of plasmid-encoded genes.

During the experiment, we noticed that Δ10 strains have several drawbacks. First, YMT53–YMT55 grew more slowly than their ancestor, S288C (Fig. 1C, SD complete) [17]. Among the Δ10 strains,YMT183 ‒ YMT185 grew even more slowly than YMT53 ‒ YMT55 (Fig. 2D, YPAD), probably because in addition to the antibiotic marker, flanking regions of the auxotrophic marker ORF were removed in YMT183‒YMT185 (Fig. 2B), which may affect transcription of neighboring genes andthereby reduce cell fitness. Second, although YMT55 exhibited normal meiosis and spore formation, YMT185 did not (data not shown), probably for the same reason mentioned above. In addition to these drawbacks, it should be noted that we have not yet confirmed whether the Δ10-based expression system enables the co-expression of 20 different genes. Further study is warranted to address these drawbacks and confirm the significance of this system in biotechnological applications.

Collectively, we developed a novel yeast-based experimental system for the establishment of cells carrying 10 different plasmids. This method holds promise as a tool to build a novel yeast cell factory.

## Author contributions

T.M. designed the project, performed most experiments, analyzed and interpreted the data, prepared the figures, and wrote the manuscript. G.D. and T.N. performed experiments, analyzed and interpreted the data, and wrote the manuscript.

## Acknowledgements

We gratefully acknowledge the financial support by Kiyofumi Yamada. This study was supported by JSPS KAKENHI grants JP23H02669 (to K.Y.) and JP22K17824 (to T.N.). G.D. would like to take this opportunity to thank the “Nagoya University Interdisciplinary Frontier Fellowship” supported by Nagoya University and JST, the establishment of university fellowships towards the creation of science technology innovation, Grand Number JPMJFS2120.

## Supplementary material

Supplemental material is available online only.

