## Supplementary figures and images for "A novel plasmid-based experimental system in *Saccharomyces cerevisiae* that enables the introduction of 10 different plasmids into cells"

### Supplemental Figure

#### Slide 1
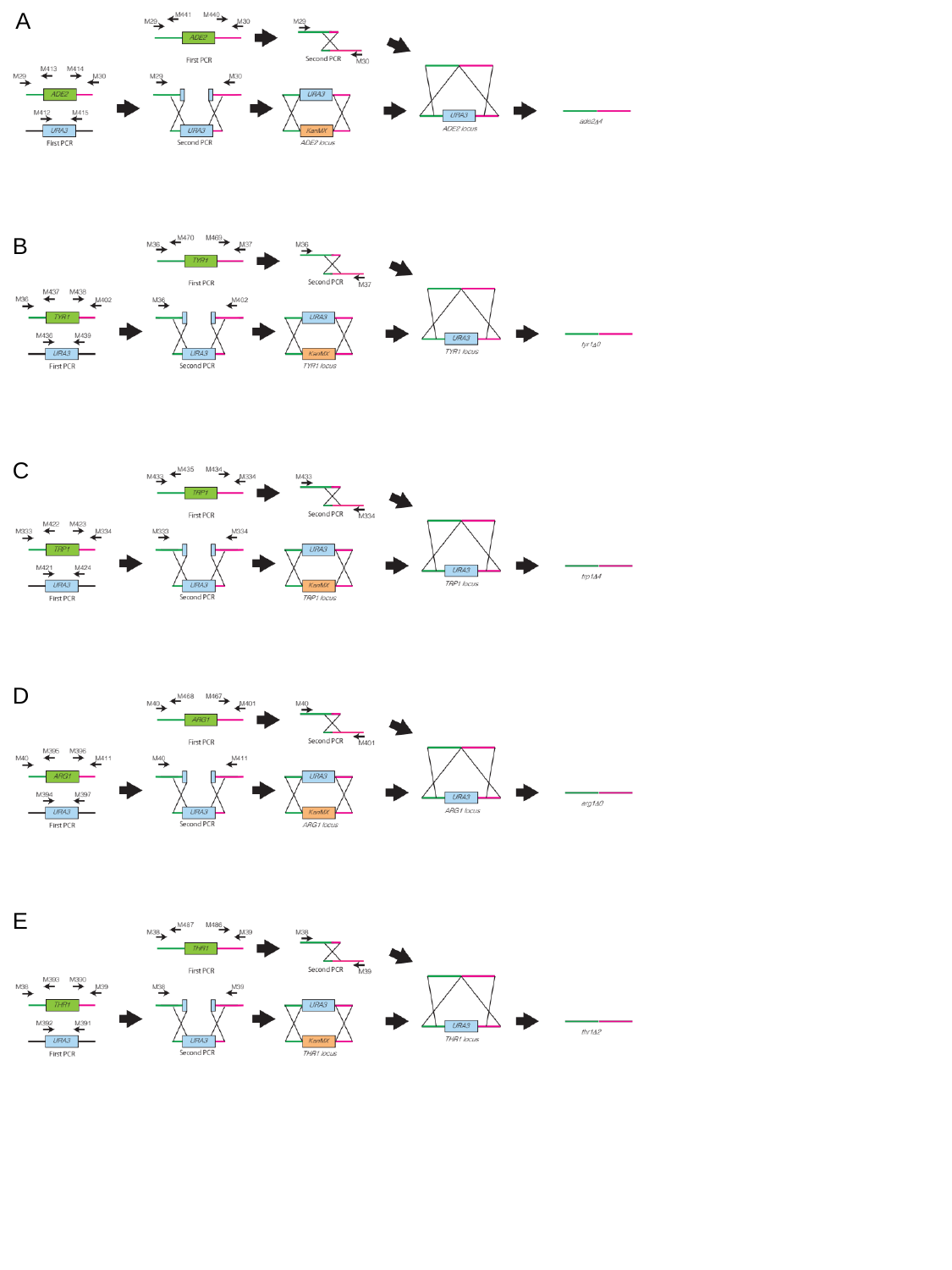

A
B
C
D
E
